# Respiratory coordination of excitability states across the human wake-sleep cycle

**DOI:** 10.1101/2025.06.03.657770

**Authors:** Andrea Sánchez Corzo, Esteban Bullón Tarrasó, Martina Saltafossi, Teresa Berther, Tobias Staudigl, Daniel S. Kluger, Thomas Schreiner

**Affiliations:** Department of Psychology, Ludwig-Maximilians-Universität München, München, Germany; Graduate School of Systemic Neurosciences, Munich, Germany; Institute for Biomagnetism and Biosignal Analysis, University of Münster, Münster, Germany; Otto Creutzfeldt Center for Behavioral and Cognitive Neuroscience, University of Münster, Münster, Germany

## Abstract

While the respiratory rhythm is increasingly recognized as a key modulator of oscillatory brain activity across the wake-sleep cycle in humans, very little is known about its influence on aperiodic brain activity during sleep. This broadband activity indicates spontaneous fluctuations in excitation-inhibition (E:I) balance across vigilance states and has recently been shown to systematically covary across the respiratory cycle during waking resting state. We used simultaneous EEG and respiratory recordings over a full night of sleep collected from N = 23 healthy participants to unravel the nested dynamics of respiration phase-locked excitability states across the wake-sleep cycle. We demonstrate a prominent phase shift in the coupling of aperiodic brain activity to respiratory rhythms as participants were transitioning from wakefulness to sleep. Moreover, respiration-brain coupling became more consistent both across and within participants, as interindividual as well as intraindividual variability systematically lessened from wakefulness and the transition to sleep towards deeper sleep stages. Our results suggest that respiration phase-locked changes in E:I balance conceivably add to sleep stage-specific neural signatures of REM and NREM sleep, highlighting the complexity of brain-body coupling during sleep.

## Introduction

A growing body of cross-species evidence continues to highlight the respiratory rhythm as a key modulator of neural processes [1]. Acting as a master clock governing perception, action, and cognition, respiration orchestrates transient states of both the brain and the body, including arousal [2,3] and cortical excitability [4,5]. Critically, this integrative brain-body link spans a wide range of time scales [6]: Fluctuations in cortical excitability, for example, covary breath-by-breath with respiration within seconds [7] but are also subject to larger-scale circadian rhythmicity over several hours [8]. While short-term neural modulation as a function of respiration phase is now relatively well understood (for review, see [9,10]), the increasingly complex, nested temporal dynamics of respiration-brain coupling remain virtually unstudied.

Respiratory modulation of human brain activity has mostly been described in the domain of neural oscillations [11,12]. With a particular focus on respiration-related changes in cortical excitability states, however, recent studies are beginning to consider the aperiodic component of neural signalling as well: This dominant broadband component of brain activity is characterised by a marked 1/f characteristic, meaning that lower frequencies carry higher power. Since the aperiodic component is hypothesized to reflect the dynamic interplay of (faster) excitatory and (slower) inhibitory currents [13], its steepness (or slope) has become an established marker of excitation-inhibition (E:I) balance in the cortex [14,15] - the larger the 1/f slope exponent, the stronger the inhibitory influence (and vice versa). Consistent results from both oscillatory and non-oscillatory (aperiodic) respiration-brain coupling [16-19] strongly suggest that spontaneous E:I dynamics in the human cortex are modulated by the breathing rhythm.

In order to go beyond existing approaches and investigate respiration-brain coupling at different time scales simultaneously, vigilance states like the wake-sleep cycle constitute a prime application of nested brain-body dynamics: While short-term respiration-locked modulations of brain activity can be observed on a cycle-by-cycle basis, longer-term changes in these modulations may be observed across different sleep stages. Here, distinct neural signatures as well as functional roles of each sleep stage in processes like homeostatic maintenance [20], neural plasticity [21], or memory consolidation [22] raise central questions as to how respiratory coupling might influence sleep characteristics.

Particularly with regard to dynamic changes in E:I balance, there is now considerable agreement that 1/f slope systematically fluctuates across vigilance and sleep stages [23-27]. At the same time, emerging studies have started to delineate the role of respiration in modulating representative neural signatures of sleep, such as slow oscillations (SOs) and sleep spindles, and their pivotal roles in memory formation and consolidation [28-30]. For example, respiratory phase-locking has been linked to the emergence and timing of coupled SO-spindle complexes, which are integral to memory reactivation during non-rapid eye movement (NREM) sleep [31].

Overall, recent findings clearly suggest that different vigilance states cannot only be distinguished by broadband, sleep stage-specific changes in aperiodic activity, but that these changes are in turn related to the generation and temporal orchestration of neural signatures of great cognitive and behavioural relevance.

Pursuing these questions, we jointly investigated respiration phase-locked modulation of excitation-inhibition balance across different time scales, quantifying aperiodic brain activity as a function of respiratory phase during wakefulness and across sleep stages. Using concurrent EEG and respiratory recordings during sleep, we provide a comprehensive characterisation of multiscale E:I coupling to the respiratory rhythm and reveal critical insights into the functionality of nested respiration-brain dynamics.

## Results

A group of N = 23 healthy volunteers were monitored for one full night in the lab, sleeping around 7 hours on average. On average, participants were awake or transitioning to NREM sleep for around 62 minutes throughout the night and spent the majority of the time of around 300 minutes in NREM sleep stages N2 and N3 (here referred to as slow-wave sleep, SWS). REM sleep was observed for around 72 minutes on average. Full sleep characteristics are provided in Table 1.

**Table 1.**
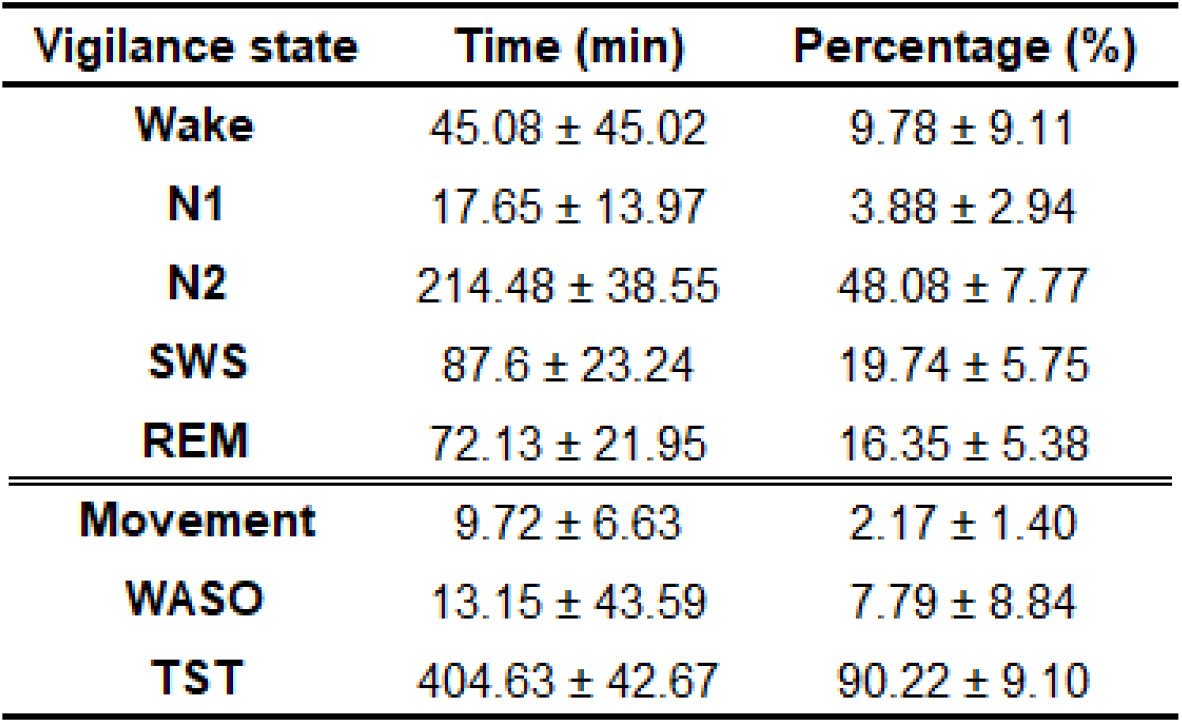
Sleep characteristics. Means ± standard deviation of time spent in each vigilance state, periods with observable movement, time spent awake after initial sleep onset, and total sleep time plus corresponding percentage relative to the full recording. N1, N2: NREM sleep stages N1 & N2, SWS: slow-wave sleep, REM: rapid eye movement sleep, Movement: movement time, WASO: wake after sleep onset, TST: total sleep time.

| Vigilance state | Time (min) | Percentage (%) |
| --- | --- | --- |
| Wake | 45.08 ± 45.02 | 9.78 ± 9.11 |
| N1 | 17.65 ± 13.97 | 3.88 ± 2.94 |
| N2 | 214.48 ± 38.55 | 48.08 ± 7.77 |
| SWS | 87.6 ± 23.24 | 19.74 ± 5.75 |
| REM | 72.13 ± 21.95 | 16.35 ± 5.38 |
| Movement | 9.72 ± 6.63 | 2.17 ± 1.40 |
| WASO | 13.15 ± 43.59 | 7.79 ± 8.84 |
| TST | 404.63 ± 42.67 | 90.22 ± 9.10 |

In order to quantify respiration phase-dependent changes in aperiodic brain activity across vigilant and sleep stages, we extracted time-resolved estimates of 1/f slope from overnight EEG recordings. Parallel respiratory recordings and sleep staging allowed us to then characterise baseline changes in neural and respiratory activity across vigilance states (i.e., wakefulness, NREM, REM sleep) as well as the coupling of these two signals during a full night of sleep (Fig. 1).

**Fig. 1.**
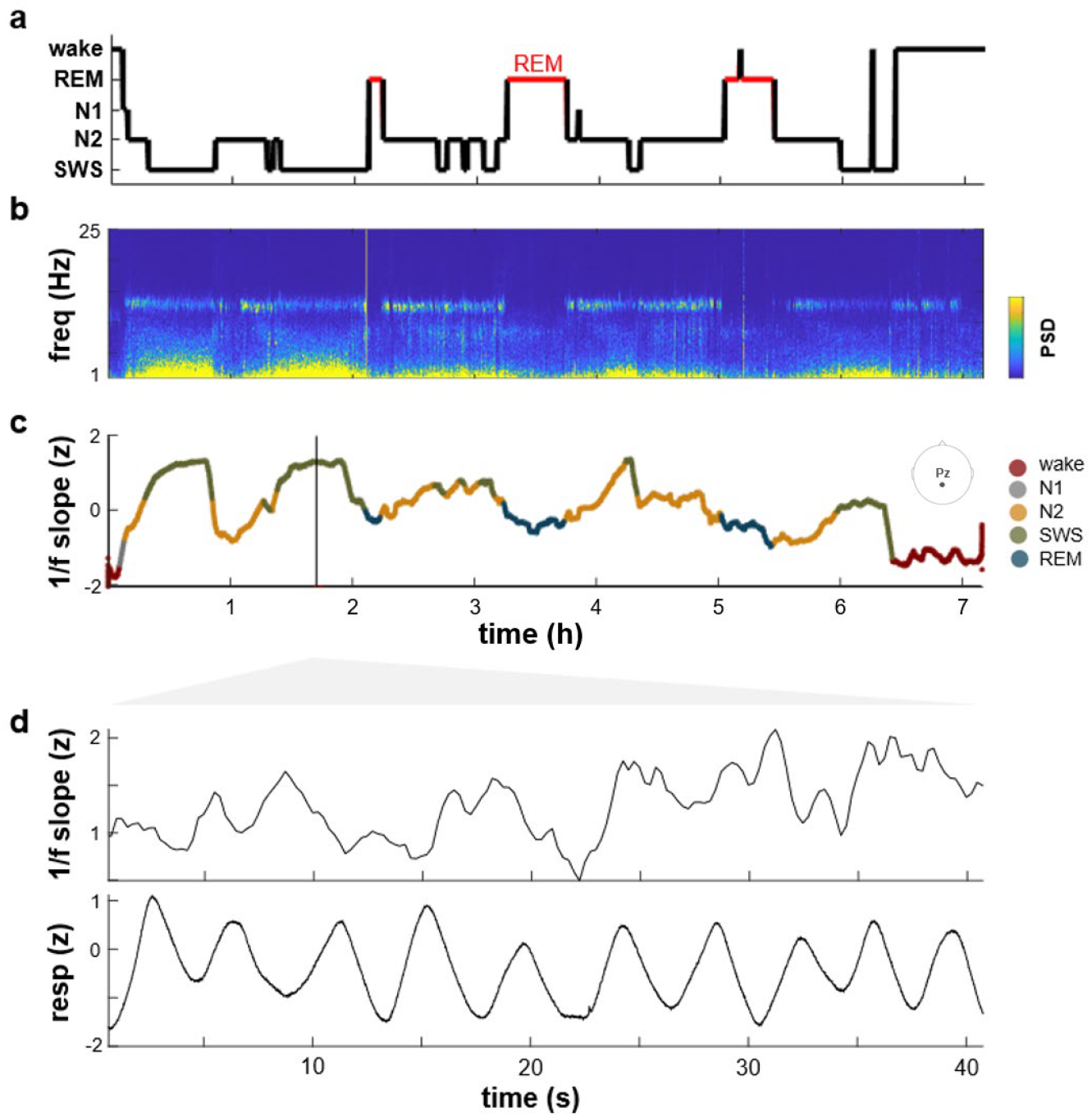
Exemplary overnight recording and methodology. **a**, Hypnogram of the full night for a single participant. **b**, Time-frequency representation (spectrogram) of the EEG data recording (exemplary channel Pz). **c**, Continuously estimated 1/f spectral slope (z-scored over the full recording) from channel Pz. Aperiodic activity was estimated within a one-second sliding window (75% overlap) applied to the whole-night time series recorded from each channel (see Methods for details). Colour code indicates the simultaneously scored sleep stage shown in panel a. The dark vertical line marks the segment enlarged below. **d**, The inset illustrates the z-scored 1/f slope (upper trace) and z-scored respiratory signal (lower trace) across an exemplary 40-second time window.

In a first analysis of *1/f slope across vigilance states*, we found aperiodic activity - averaged across channels within each participant - to be significantly modulated by polysomnographic stage (Friedman test: χ^2^(4) = 88.24, *p* < .001). Pairwise comparisons across stages revealed lower overall 1/f slope during wakefulness compared to N2 (mean rank difference *M* = -2.91, *p* < .001), SWS (*M* = -3.91, *p* < .001), and REM (*M* = -1.78, *p* = .001; Tukey-Kramer-corrected), indicating the greatest influence of excitatory currents during wakefulness (Fig. 2a). Moreover, 1/f slope during the transition from wake to sleep (i.e., during sleep stage N1) was significantly lower than during N2 (*M* = -1.96, *p* < .001) and during SWS (*M* = -2.96, *p* < .001), suggesting similar excitatory dynamics during wakefulness and the transition to NREM sleep onset (N1). Finally, the overall highest 1/f slope during SWS, indicative of strong inhibitory currents, was also significantly greater than aperiodic activity during REM (*M* = 2.13, *p* < .001).

**Fig. 2.**
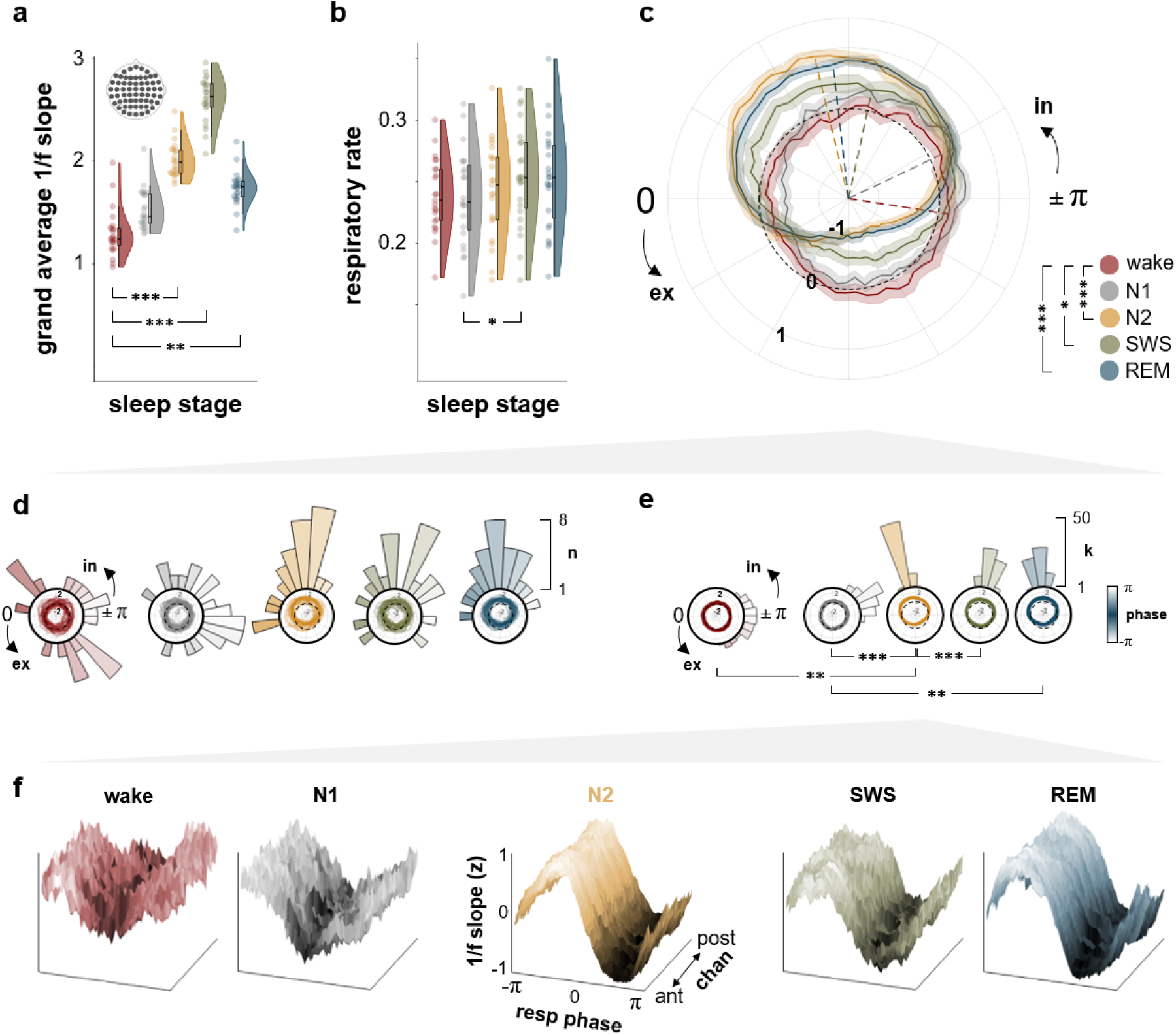
Sleep stage-dependent changes in aperiodic activity and its coupling to respiration phase. **a**, Group-level distribution of 1/f slope across polysomnographic stages, averaged across all k = 60 EEG channels and irrespective of respiration phase. Tukey-Kramer-corrected significance is shown relative to wakefulness; full statistical results are provided in the main text. **b**, Group-level distribution of respiratory rates across stages. Across all pairwise comparisons, only the difference between N1 and SWS was significant (*p* = .041, Tukey-Kramer-corrected). **c**, Polar representation of group-level mean 1/f slope ± SEM across respiration phase per polysomnographic stage, averaged across all k = 60 channels. Coloured dashed lines indicate mean directions of the respective circular distributions. FDR-corrected significance is shown relative to wakefulness; full statistics on differences in mean direction are provided in the main text. **d**, Polar histograms show the circular distribution of mean directions in 1/f slope across the n = 23 participants per stage. Polar plots show individual courses of aperiodic activity over respiration phase. FDR-corrected tests against circular uniformity were highly significant for all polysomnographic stages. Statistical results are provided in the main text. **e**, Polar histograms show the circular distribution of group-level mean directions in 1/f slope across the k = 60 electrodes per stage. Polar plots show electrode-specific courses of aperiodic activity over respiration phase. FDR-corrected tests against circular uniformity were highly significant for all sleep stages. Statistical results are provided in the main text. **f**, Detailed visualisation of topographic consistency in group-level average 1/f slope across all electrodes, sorted from anterior to posterior location.

In light of these sleep stage-dependent changes in aperiodic activity, we next investigated baseline *changes in respiratory rate across sleep stages* before conducting joint analyses of the two dimensions. A Friedman test showed significant variation in respiratory rate across sleep stages (χ^2^(4) = 10.54, *p* = .032), driven by a significant increase in respiration rate from N1 to SWS (*M* = 1.30, *p* = .041), the stages with the lowest and highest overall respiration rates, respectively (Fig. 2b).

Since our main focus was on respiration-phase dependent modulation of aperiodic activity across stages of the wake-sleep cycle, we extracted 1/f slope estimates per participant, EEG channel, polysomnographic stage, and respiration phase. To this end, and in keeping with previous approaches [16,18,19], continuous respiration phase vectors were segmented into n = 60 phase bins (see Methods). From scalp-average courses of *1/f slope across the entire respiration cycle*, we computed individual circular means per stage using the *circstat* toolbox for Matlab [32]. We then conducted pairwise statistical comparisons to quantify the likelihood that circular means of slope ~ phase of any two polysomnographic stages were sampled from the same distribution. Pairwise Watson’s U^2^ tests revealed that distribution of 1/f slope over respiration phase during wakefulness (mean angular direction α= 2.99 or 171.09°) was significantly different from those during N2 (α= -1.33 or -76.43°, U^2^ = 0.41, FDR-corrected *p* < .001), SWS (α= -1.78 or -102.14°, U^2^ = 0.17, *p* = .031), and REM (α= -1.45 or -83.33°, U^2^ = 0.34, *p* < .001; Fig. 2c). While wakefulness was characterised by decreased aperiodic activity during late inspiration and increased 1/f slope during late expiration, this pattern was notably reversed for all subsequent sleep stages, starting with N2. During the transition to NREM sleep (i.e., during N1), the distribution of aperiodic activity across respiration phase was similar to wakefulness (α= -2.70 or -154.60°), but significantly different from N2 (U^2^ = 0.41, *p* = .022) and REM (U^2^ = 0.28, *p* = .022). Finally, significantly different distributions were found for SWS compared to N2 (U^2^ = 0.18, *p* = .024) and REM (U^2^ = 0.09, *p* = .031).

In summary, the later sleep stages N2, SWS, and REM showed a marked increase in aperiodic activity - associated with increased inhibition - during the entire inspiratory phase, while 1/f slope was decreased (indicating *decreased* inhibition) during expiration. Given that respiration-brain coupling typically varies greatly across participants, we sought to quantify *interpersonal variability* in our findings. Polar histograms in Fig. 2d show the prevalence of mean angular directions for slope ~ phase in our sample, separately for each sleep stage. While individual distributions of aperiodic activity over respiration phase were quite variant during wakefulness and light sleep (circular variance: s_wake_ = 38.46° and s_N1_ = 25.28°, respectively), deeper sleep stages like N2 (s_N2_ = 10.31°) and REM (s_REM_ = 10.08°) showed less interpersonal variability. In other words, not only did we observe a shift in respiratory phase coupling with transient excitability states during N2 and REM (compared to wakefulness and N1), but these changes in phase-dependent aperiodic activity also were more consistent across participants.

As a final dimension of interest, we investigated *topographic variability*, i.e. how consistently we would observe vigilant and sleep stage-dependent changes in respiration-brain coupling across the scalp. To this end, we extended our analysis of individual circular means to phase-resolved estimates of 1/f slope based on single-channel EEG data. For each participant and polysomnographic stage, we computed the circular variance of mean angular directions across all k = 60 channels. A Friedman test showed significant variability differences between stages (χ^2^(4) = 36.42, *p* < .001). Lowest cross-channel variability was found during N2 periods (circular variance: s_N2_ = 5.43°) at which variance was significantly lower than during wakefulness (s_wake_ = 18.27°; *M* = -1.78, *p* = .001), N1 (s_N1_ = 23.26; *M* = -2.57, *p* < .001), and SWS (s_SWS_ = 16.01; *M* = -1.91, *p* < .001). Finally, variability in mean angular directions across channels was lower during N1 compared to REM (s_REM_ = 9.35°; *M* = -1.65, *p* = .004; Fig. 2e).

Overall, our results manifest respiration as a critical modulator of brain activity across brain-body states. We demonstrate a prominent phase shift in the coupling of aperiodic brain activity to respiratory rhythms as participants were transitioning from wakefulness to sleep. Moreover, respiration-brain coupling became more consistent both across and within participants (Fig. 2f), as interindividual as well as intraindividual variability systematically lessened from wakefulness and early sleep onset (N1) towards deeper sleep stages.

## Discussion

The main aim of this work was to unravel the nested dynamics of respiration phase-locked excitability states across the wake-sleep cycle. Across polysomnographic stages, we found that overall aperiodic activity increased from wakefulness/N1 towards deeper NREM sleep stages N2 and SWS as well as REM sleep. These results are consistent with previous findings using the same frequency range for slope estimation (1-40 Hz; [24,33]) and represent a continuous increase in cortical inhibition states for later sleep stages compared to wakefulness. While we also report slightly increased respiratory rate in line with earlier studies [34-36], changes in breathing frequency clearly cannot account for these baseline differences in aperiodic activity. This corroborates previous comprehensive correlation analyses [18] in which we found no meaningful link between individual respiratory rate and 1/f slope across multiple data sets.

Our analysis of respiration phase-locked excitability changes across polysomnographic stages revealed that the coupling between E:I dynamics and respiration phase was by no means uniform across the wake-sleep cycle, but showed a systematic phase shift from wakefulness/N1 to deeper NREM (N2, SWS) and REM stages: During wakefulness, aperiodic activity was increased around the exhalation-to-inhalation transition (associated with higher inhibition) and decreased around the inhalation-to-exhalation transition (associated with higher excitation), which closely follows the observations of our previous work [18]. During both NREM and REM sleep, this phase locking was significantly altered, with steeper 1/f slope (i.e., stronger inhibitory activity) during inspiration and flattened 1/f slope (stronger excitation) during exhalation. Notably, these phase shifts were increasingly consistent both between and within participants.

On a general level, these results provide further evidence that the phase of the respiratory rhythm impacts neural activity during both wakefulness and sleep, underscoring the complexity of brain-body states [6]. To date, the impact of respiration on both oscillatory and aperiodic activity has been better described for wakefulness [11,17,37], with the focus increasingly on modulations of excitability states and E:I balance [16,18]. These respiration-locked changes in excitability states are not merely accidental, but have been shown to actually facilitate perception [16], which raises the question whether equivalent phase-dependent E:I modulation during sleep serves an equally functionally relevant purpose. Indeed, our observation of differential respiratory phase locking for wakefulness vs NREM/REM sleep hints at different functional roles for respiration in the organisation of excitability across vigilance states. Moreover, recent rodent work by Jung and colleagues [38] has demonstrated that the preferred timing of respiration-entrained discharges in the mouse parietal cortex varied between vigilance states, which suggests distinct, state-specific underlying mechanisms of respiration-brain coupling across the wake-sleep cycle.

To begin with, the potential role of respiration in modulating REM-related brain activity is entirely unknown in humans. Leaving respiratory interactions aside, the differential microarchitecture of tonic and phasic REM sleep has been related to changes in aperiodic neural activity [25]. Here, E:I balance would be leaning towards inhibitory activity during tonic REM sleep and towards higher excitation during phasic REM sleep, respectively [39]. The extent to which these baseline dynamics between quiescent and more active states may, in turn, covary with the respiratory rhythm in humans, however, is unclear and opens promising avenues for future investigation. A first indication of such coupling differences between phasic and tonic REM sleep comes from rodent work by Hammer and colleagues [40]: The authors showed that cross-frequency coupling of theta and gamma oscillations in mice was differentially related to the respiratory rhythm during phasic vs tonic REM sleep. Of note, theta-gamma coupling overall is strongest during REM sleep [41] and thus thought to be of critical importance for state-specific information processing in the rodent cortex [42]. Hence, it appears that respiratory characteristics like phase and frequency should be taken into account when trying to understand the complexities of distinct sleep stage signatures and their functional relevance.

With regard to NREM sleep, there is now growing evidence that the respiratory rhythm impacts NREM-specific oscillatory signatures [28,29,43,44]. Here, the precise phase locking of cortical slow oscillations (SOs) and thalamo-cortical sleep spindles in particular provides a common explanatory reference frame for systematic comodulation of both oscillatory and aperiodic activity during NREM sleep: Slow oscillations reflect low-frequency fluctuations (<1 Hz) of membrane potentials, with SO downstates being indicative of hyperpolarization whereas upstates are related to depolarization [45]. Consequently, SOs drive temporal windows of excitation-inhibition dynamics not only in cortical but also in subcortical areas [46]. We have previously shown that SO downstates, i.e. a slow oscillatory indicator of neural silence and inhibition, preferentially emerged before the inhalation peak [47]. In contrast, sleep spindles - faster (12-16 Hz) thalamic oscillations known to systematically lock to excitatory SO upstates - preferentially emerged during early exhalation. These distinct preferred phases of oscillatory signatures of inhibitory and excitatory states closely resemble the phase locking pattern we observed here for aperiodic activity: During wakefulness, the distribution of 1/f slope across respiration phase is such that exhalation is related to higher inhibition, which is in line with previous findings [18]. After sleep onset, the prominent phase shift in respiratory coupling we describe above rearranges the association of E:I balance and respiration phase, so that inhibition is now lower during the expiration phase. Conceivably, this increase in baseline excitability now allows excitatory SO upstates and nested sleep spindles to emerge during expiration, as previously observed.

Although further work is of course needed to explore this potential mechanism in more detail, we provide first evidence that respiration phase-locked changes in aperiodic activity contribute to the emergence of SO-spindle complexes at a preferred respiratory phase. Critically, this does not mean that SO upstates will emerge automatically, e.g. during REM sleep whose E:I dynamics were similarly coupled to respiratory phase. Rather, it seems likely that respiratory modulation of aperiodic activity lays the necessary foundation for sleep stage-specific neural signatures to occur in phase with the breathing cycle. In case of NREM sleep, SO-spindle complexes play a crucial role in memory consolidation [48,49] and the extent of their coupling to respiration phase is predictive of memory performance [47].

Taken together, our results suggest that respiration orchestrates these functionally and behaviourally relevant neural signatures through different mechanisms. Respiration phase-dependent changes in aperiodic activity across the wake-sleep cycle are clearly observable, increasingly consistent as participants transition from wakefulness to sleep, and highly coherent across the cortex. With regard to functional relevance, these changes in E:I balance conceivably add to the respective, sleep stage-specific neural signatures of REM and NREM sleep, highlighting the complexity of brain-body coupling during sleep.

## Materials and Methods

### Participants

Twenty-three volunteers (15 female, age 24.09 ± 3.51 [mean ± SD]) participated in the study. All participants reported having no respiratory or neurological disease (including sleep apnea) and gave written informed consent prior to all experimental procedures. Pre-study screening involved several questionnaires, including the Pittsburgh Sleep Quality Index (PSQI; [50]), the Morningness-Eveningness Questionnaire [51], and a custom questionnaire assessing general health and stimulant use. Results indicated that none of the participants were taking any medication at the time of the experimental session, and all were free from neurological and psychiatric disorders. All participants reported generally good sleep quality. Furthermore, they had not been on a night shift for at least 8 weeks before the experiment. All participants were instructed to avoid alcohol the evening before and caffeine on the day of the experimental sessions. They received course credit or financial reimbursement in return for their participation. The study was approved by the ethics committee of the Department of Psychology, Ludwig-Maximilians-Universität München).

### Procedure

On experimental days, participants arrived at the sleep laboratory at around 8:30 p.m. The experimental sessions started with the setup for polysomnographic recordings during which electrodes for electroencephalographic (EEG), electrooculographic (EOG), and electrocardiographic (ECG) recordings were applied. In addition, a thermistor airflow sensor was attached to record respiration. Several days before the experimental session, participants were habituated to the environment by having an adaptation nap in the sleep laboratory. The data presented here was collected as part of a broader project investigating sleep and memory. Participants completed an associative memory task before and after sleep; however, these behavioral data are beyond the scope of the current work and will be reported elsewhere. The sleep period began at around 11:30 p.m. and all participants slept for around 7-8 hours (for sleep characteristics, see Table 1). In the morning, participants were awoken from light sleep (sleep stage N1 or N2).

### EEG and respiratory recordings

An EEGo 65-channel EEG system (ANT Neuro, Enschede, Netherlands) was used to record the EEG throughout the experiment. Impedances were kept below 20 kΩ. EEG signals were referenced online to electrode CPz and sampled at a rate of 512 Hz. Respiration was recorded in parallel using an EMBLA thermistor airflow sensor.

### EEG preprocessing and sleep staging

All EEG and respiratory data preprocessing was done in Fieldtrip for Matlab [52]. EEG data were resampled to 100 Hz and visually inspected for data loss at the very end of the recording (i.e., during waking up). Power line artefacts were removed using a discrete Fourier transform (DFT) filter on the line frequency of 50 Hz and its harmonics (including spectrum interpolation). Faulty channels were identified using kurtosis and variance criteria and interpolated with spline interpolation, if needed (2.13 ± 1.06 channels interpolated on average; M ± SD). We subsequently applied independent component analysis (ICA) on the filtered data to remove artificial components (0.52 ± 1.34 removed on average) and re-referenced all channels to the common average. Sleep staging was carried out offline according to standard criteria [53] by two independent raters.

### Respiratory preprocessing and phase extraction

After resampling to 100 Hz, transition points of inspiration-to-expiration (i.e., peaks in the respiratory time series) and expiration-to-inspiration (troughs) were identified in the respiration time course. Due to the great variability in respiration depth across the hours-long recording, peaks and troughs were identified using a 10s moving window applied in 5s increments. Within each window, the respiratory signal was normalised and Matlab’s *findpeaks* function detected peaks with a minimum peak-to-peak distance of 2s and a minimum peak prominence of *z* = 0.1. After removal of duplicate peaks, troughs were identified as local minima between two respiration peaks. Finally, phase angles were linearly interpolated from trough to peak (−π to 0) and peak to trough (0 to π) to yield respiration cycles centred around peak inspiration (i.e., phase 0).

### Computation of time-resolved aperiodic activity

Single-sensor time series of the entire recording within this ROI were entered into the SPRiNT algorithm [54] with default parameter settings and subsequently averaged. SPRiNT is an extension of the *specparam* algorithm [55] and uses a short-time Fourier transform (frequency range 1-40 Hz) to estimate the aperiodic component in neural time series within a moving window (1s width, 75% overlap).

Thus, SPRiNT yielded time series of the aperiodic component (i.e., the 1/f exponent) with a temporal resolution of 250 ms and a frequency resolution of 1 Hz. In order to relate 1/f slope time series to the respiratory signal, we extracted respiratory phase at all time points for which slope had been fitted (i.e., the centres of each moving window). In keeping with previous work [16,18,19,56], the respiratory cycle was binned into n = 60 equidistant, overlapping phase bins. Moving along the respiration cycle in increments of Δω = π/30, we collected 1/f slope fits computed at a respiration angle of ω ± π/10. On these estimates, we then computed bin-wise averages of 1/f slope, yielding quasi-continuous ‘phase courses’ of the aperiodic component.

#### Statistical analyses

#### Aperiodic activity across sleep stages

To determine overall sleep stage-dependent differences in aperiodic activity, we computed individual mean 1/f slopes per sleep stage, averaged across respiration phases and EEG channels. We used a non-parametric Friedman test to determine significant group-level slope changes across sleep stages with subsequent Tukey-Kramer correction using Matlab’s *multcompare* function (see Fig. 2a).

#### Descriptive analyses

To compute individual respiratory rates per sleep stage, we counted inspiratory peaks per minute over time and calculated the average. To identify these peaks in the normalized respiration signal, we applied Matlab’s findpeaks function, setting the minimum peak prominence to z = 0.5 (Fig. 2b). Based on sleep staging, we computed the average time spent in each sleep stage, the average amount of time during which participants were moving, as well as the average duration of wakefulness after sleep onset. Descriptive sleep characteristics are provided in Table 1.

#### Aperiodic activity as a function of respiration phase across sleep stages

We used individual matrices of 1/f slope estimates across 5 sleep stages X 60 respiration phase bins (again, averaged all channels) to investigate sleep stage-dependent differences in the coupling of aperiodic activity to respiration phase. Group-level mean courses of slope ~ phase per sleep stage were Rayleigh-tested against uniformity using the *circstat* toolbox for Matlab [32] and subsequently FDR-corrected (see Fig. 2c). Sleep stage-dependent shifts in the mean directions of slope ~ phase distributions were quantified with pairwise Watson’s U^2^ tests using the *watsons_U2* function for Matlab ([56]; see Results). Finally, polar histograms describe the consistency of our findings both across participants and EEG channels (Fig. 2d-f).

## Acknowledgements

Andrea Sánchez Corzo is supported by a Marie Skłodowska-Curie Actions Postdoctoral Fellowship (grant agreement number 101151627) within the framework of the HORIZON-MSCA-2023-PF-01-01 call. Tobias Staudigl is supported by an ERC Starting Grant (grant agreement number 802681) awarded by the European Research Council. Daniel Kluger is supported by the Innovative Medical Research (IMF) programme by the University of Münster (grant ID KL 1 2 22 01) and the German Research Council (DFG; grant agreement number KL 3580/1-1). Thomas Schreiner is supported by an Emmy-Noether grant (grant agreement number 492835154) awarded by the German Research Council (DFG).

